# Replication competition drives the selective mtDNA inheritance in *Drosophila* ovary

**DOI:** 10.1101/2025.04.29.650984

**Authors:** Cheng Zhang, Zhe Chen, Hansong Ma, Hong Xu

## Abstract

Purifying selection that limits the transmission of harmful mitochondrial DNA (mtDNA) mutations has been observed in both human and animal models. Yet the precise mechanism underlying this process remains undefined. Here, we present a highly specific and efficient *in situ* imaging method capable of visualizing mtDNA variants that differ by only a few nucleotides at single-molecule resolution in *Drosophila* ovaries. Using this method, we revealed that selection primarily occurs within a narrow developmental window during germline cysts differentiation. At this stage, the proportion of the deleterious mtDNA variant decreases without a reduction in its absolute copy number. Instead, the healthier mtDNA variant replicates more frequently, thereby outcompeting the co-existing deleterious variant. These findings provide direct evidence that mtDNA selection is driven by replication competition rather than active elimination processes, shedding light on a fundamental yet previously unresolved mechanism governing mitochondrial genome transmission.

## INTRODUCTION

The mitochondrial genome (mtDNA) is prone to accumulating mutations due to its proximity to damaging free radicals and the lack of canonical DNA repair mechanisms (Taylor & Turnbull, 2005). Yet, the infrequency of deleterious mtDNA mutations in populations underscores the presence of effective mechanisms limiting their transmission across generations (Rand, 2008). In nearly all animal species, mtDNA is exclusively transmitted through the maternal lineage (Wallace, 2008). During oogenesis, the genome goes through a genetic bottleneck that promotes random segregation and rapid genetic drift of mitochondrial variants (Cree *et al*, 2008; Jenuth *et al*, 1996). Oocytes with excessive mutations would, in principle, suffer from compromised energy metabolism and be eliminated during germ cell development. However, the frequency of spontaneous mtDNA mutations, estimated in the range of 10^-5^-10^-6^ (Pesole *et al*, 1999), is too low for an effective selection at the cellular level. In recent years, purifying selection, where the frequency of deleterious mutations even at low level is further reduced in female germline, has been observed in human and various animal models (Fan *et al*, 2008; Hill *et al*, 2014; Lieber *et al*, 2019; Ma *et al*, 2014; Stewart *et al*, 2008; Wei *et al*, 2019). A clear reduction of mutation load on mtDNA occurs during human primordial germ cell specification and migration (Floros *et al*, 2018), and germline selection is considered a major force shaping mtDNA variant characteristics in human populations (Wei *et al*., 2019). In mouse models, strong purifying selection against pathogenic mtDNA mutations, presumably at the organelle level, occurs during folliculogenesis (Ru *et al*, 2024). Nonetheless, the cellular mechanisms underlying these selections are unclear.

The selective inheritance against deleterious mtDNA mutations has also been demonstrated in *Drosophil*a *melanogaster* (Hill *et al*., 2014; Lieber *et al*., 2019; Ma *et al*., 2014), allowing the detailed dissection of mtDNA selection in this genetically tractable model organism (Chen *et al*, 2020; Chiang *et al*, 2019; Palozzi *et al*, 2022). Previous studies have roughly mapped the timing of mtDNA selection to a developmental window spanning the late germarium to early stages of egg chambers (Hill *et al*., 2014; Ma *et al*., 2014). *Drosophila* oogenesis begins with the asymmetric division of a germline stem cell (GSC) at the anterior tip of a germarium, producing a cystoblast that undergoes four rounds of division with incomplete cytokinesis, forming a 16-cell cyst in germarium region 2A (**Figure 1A)**. Moving posteriorly, the 16-cell cysts will be encased and compressed by migrating somatic cells into the characteristic lens shape in the 2B region (**Figure 1A)**. At this stage, germ cells begin to differentiate into one future oocyte and 15 nurse cells. Once fully enveloped by follicle cells, the cyst buds off from the posterior end of the germarium and continues the development as an egg chamber (Bastock & St Johnston, 2008; de Cuevas *et al*, 1997; Hinnant *et al*, 2020; Spradling, 1993).

**Figure 1.**
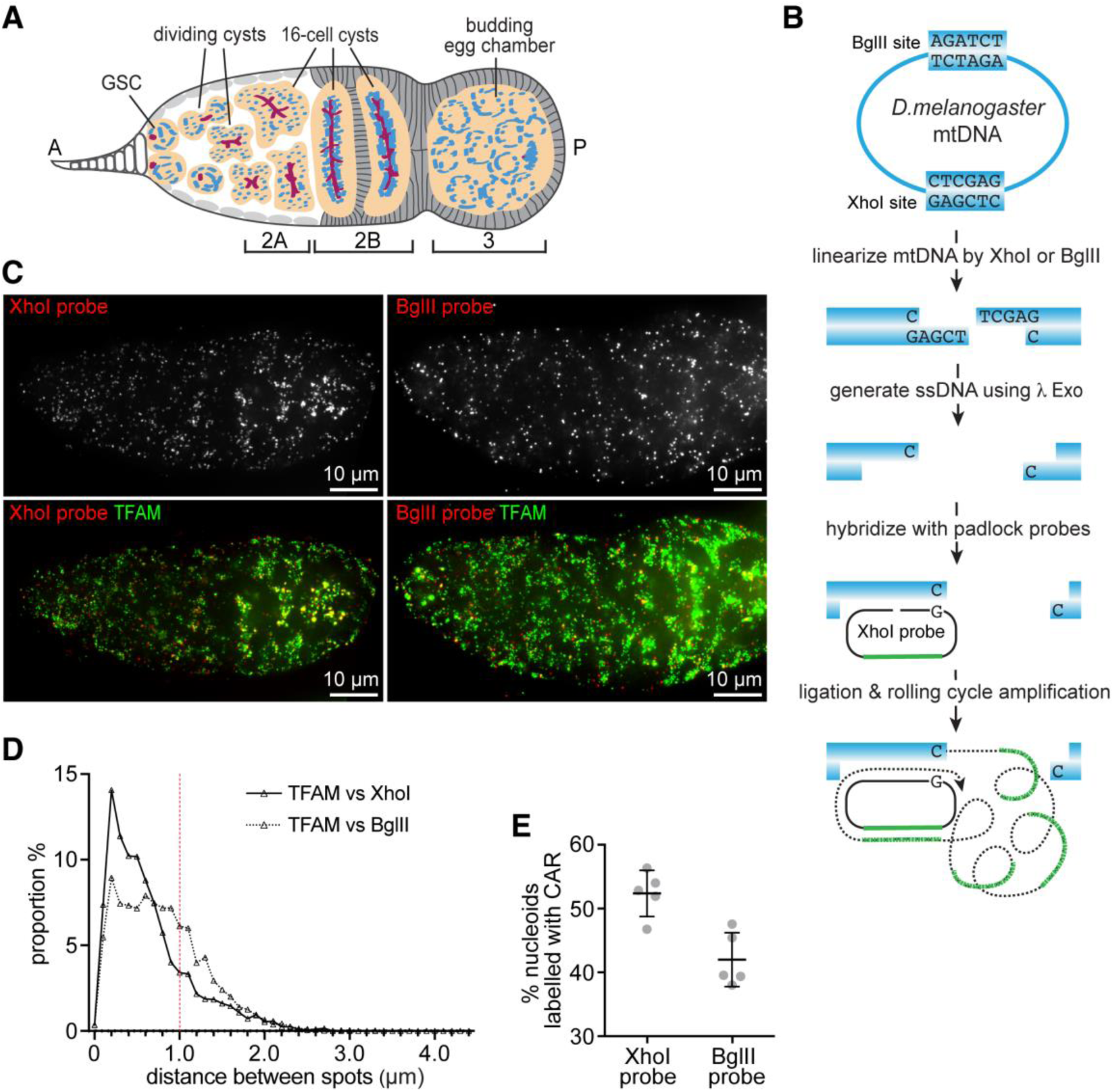
CAR assay effectively detects mitochondrial nucleoids in the *Drosophila* germarium. Schematic representation of the CAR assay for detecting mtDNA *in situ*. (**A**) Diagram of a *Drosophila* germarium illustrating successive developmental stages from anterior (A) to posterior (P). Germline stem cells (GSCs), dividing cysts (DC), regions 2A and 2B of 16-cell cysts, budding egg chambers, mitochondria (blue) and fusome (red) are depicted. (**B**) Scheme of CAR assay. The wild-type *D. melanogaster* mitochondrial genome contains a single BglII and a single XhoI site. Endogenous mtDNA is digested by XhoI or BglII restriction enzyme in tissue, followed by Lambda exonuclease (λ Exo) treatment to generate single-stranded DNA (ssDNA) with a 3’-OH group. The padlock probes are designed with two key components: a DNA sequence complementary to the ssDNA adjacent to XhoI or BglII restriction site and a unique sequence tag for subsequent detection. Upon annealing, circular ligation, and rolling circle amplification (RCA), a long, periodic ssDNA product covalently linked to the mtDNA molecule at the restriction enzyme site is synthesized. The CAR products are visualized by fluorescence *in situ* hybridization using probes specific to the unique sequence tags on each padlock probe. (**C**) CAR signals detected using XhoI (left) or BglII (right) probes colocalize with TFAM-labeled mitochondrial nucleoids in the germarium. *Drosophila* ovaries endogenously expressing TFAM-mNeonGreen were hybridized with either XhoI (left) or BglII (right) probes. Images are projections of 10 z-stacks (0.3 μm/stack). Scale bar, 10 μm. (**D**) Frequency distribution plots of distances between TFAM and CAR signals. The shortest distances between the centers of TFAM and CAR puncta were measured (XhoI probe, n=3789; BglII probe, n=4147). The average puncta diameter (∼1 μm) is indicated by a dashed line. Bin size, 0.1 μm. (**E**) Quantification of CAR assay efficiency in detecting mitochondrial nucleoids. The percentage of CAR signals relative to the total number of nucleoids in a germarium is shown. Each data point represents one germarium (n = 5). Data are presented as mean± SD.

In the germarium, a series of developmentally orchestrated mitochondrial processes that ensure the transmission of healthy mtDNA have been revealed (Chen *et al*., 2020; Palozzi *et al*., 2022). In dividing germline cysts, mtDNA transcription is largely inactive (Wang *et al*, 2019). mtDNA replication, which depends on transcription for initiation (Liu *et al*, 2022), is also quiescent (Hill *et al*., 2014). Meanwhile, mitochondria undergo active fission at this stage (Chen *et al*., 2020; Lieber *et al*., 2019), which, together with the lack of mtDNA replication, promotes mtDNA segregation. By region 2A, each mitochondrion in 16-cell cysts contains, on average, a single mtDNA nucleoid (Chen *et al*., 2020). This links the function of individual mitochondria to the quality of genome they carry, allowing potential mtDNA selection at the organelle level. From GSCs to rounded 16-cell cysts in region 2A, mitochondrial respiration is inactive (Wang *et al*., 2019), presumably due to the lack of mtDNA transcription that generates core components of electron transport chain complexes. In region 2B, the mechanical stress from surrounding follicle cells activates mitochondrial respiration through a Myc-mediated transcriptional boost in differentiating 16-cell cysts (Wang *et al*, 2023). Interestingly, mtDNA replication commences concurrently with, and appears depending on the activation of mitochondrial respiration, as mtDNA replication is severely impaired in region 2B cysts carrying homoplasmic *mt:CoI^T300I^*, a temperature sensitive lethal mtDNA mutation that severely disrupts cytochrome C oxidase (Hill *et al*., 2014). These observations have led to a proposal that wild type mtDNA or healthy genomes would be replicated more frequently and thereby outcompete deleterious variants (Chen et al, 2020). In support of this idea, severe inhibition of mtDNA replication diminishes mtDNA selection (Zhang *et al*, 2019). While this model of replication competition is logically compelling, it has yet been empirically demonstrated that deleterious mtDNA variants indeed replicate less than wild-type genome in the same germ cell. Additionally, a developmentally regulated mitophagy - programmed germline mitophagy (PGM) that begins at region 2A as germ cells enter meiosis (Palozzi *et al*., 2022), has also been shown to contribute to the selective inheritance (Lieber *et al*., 2019; Palozzi *et al*., 2022). However, given that PGM occurs regardless of the presence of mtDNA mutations (Palozzi *et al*., 2022), it is unclear whether PGM impacts mtDNA selection directly by eliminating defective mitochondria or indirectly through other processes such as mitochondrial protein turnover. Hence, the underlying mechanism of selective inheritance remains unsettled. Differentially visualizing co-existing mitochondrial genomes in ovaries is necessary to understand the timing and mechanisms of mtDNA selection.

Currently, most mtDNA variants in *D. melanogaster* were generated using a selection scheme based on mitochondrially targeted restriction enzymes (Xu, 2008). These variants differ from the wild-type genome on merely a single nucleotide or small indel (insertion-deletion) on the corresponding enzyme sites (Xu, 2008). Distinguishing single nucleotide polymorphisms (SNPs) on mtDNA *in situ* remains a technical challenge. Previous studies generated heteroplasmic fly models containing *mt:CoI^T300I^* from *D. melanogaster* and wild-type *Drosophila yakuba* mtDNA (Ma & O’Farrell, 2015). Mitochondrial genomes from these two sibling species share ∼93% sequence identity in coding regions, but the non-coding AT regions, which contain the origins of replication, are highly divergent (Clary & Wolstenholme, 1985). These polymorphisms enable the design of a pool of genome-specific probes required for single-molecule fluorescence *in situ* hybridization (smFISH), as well as a more advanced method recently developed termed smFISH-HCR, to visualize mtDNA and thereby study the underlying mechanisms of mtDNA selection (Chandrasegaram *et al*, 2025; Lieber *et al*., 2019). By providing functional cytochrome c oxidase activities, the *D. yakuba* mtDNA can be maintained at lower percentage with *mt:CoI^T300I^* at the restrictive temperature for many generations (Ma & O’Farrell 2016). However, it is quickly outcompeted by wild-type *D. melanogaster* mtDNA in the *D. melanogaster* nuclear background (Ma & O’Farrell, 2015). Hence, the potential mito-nuclear incompatibility associated with the *D. mel/yak* heteroplasmic line complicates the selective pressures on mtDNA and raises concerns about the validity and physiological relevance of this interspecies heteroplasmic model for studying mtDNA selective inheritance. It is preferable to use heteroplasmic variants from the same species to minimize confounding factors for understanding mechanisms of mtDNA transmission.

In this study, we introduce a highly specific and efficient *in situ* imaging method that enables the visualization of mtDNA genomes differing by only two loci in tissues. Using this approach, we reveal a selective replication advantage of the healthy mtDNA variant in differentiating cysts, where it outcompetes the co-existing deleterious variant. This precisely timed developmental window acts as a mitochondrial quality control checkpoint, increasing healthy mtDNA levels before the onset of large-scale mitochondrial biogenesis in developing egg chambers. Our findings provide key insights into the mechanisms driving mtDNA selection and inheritance, establishing a powerful framework for future studies on mtDNA dynamics with spatial resolutions across different tissues and model systems.

## RESULTS

### CAR assay effectively and specifically detects mtDNA SNPs *in situ*

To distinguish mtDNA variants with single nucleotide polymorphisms, we first tested a CRISPR-Cas9-mediated proximity ligation (CasPLA) assay in *Drosophila* ovaries (Zhang *et al*, 2018). However, the resulting signals were diffused within the mitochondrial network (**Figure S1**), lacking the spatial resolution needed to differentiate mtDNA genotypes in heteroplasmic ovaries. Currently, most mtDNA variants in *D. melanogaster* are generated using mitochondrially targeted restriction enzymes, which select for mutations that abolish enzymes’ cleavage sites (Xu, 2008). Taking advantage of this molecular nature, we improved a target-primed rolling circle amplification (RCA) method (Larsson *et al*, 2004) to distinguish genomes that differ by two loci with high efficiency and specificity (**Figure 1B, Figure S2A**). In our method, referred to as “CAR” (<u>C</u>ircular DNA <u>A</u>mplification at <u>R</u>estriction enzyme sites), endogenous mtDNA was digested with restriction enzymes and treated with Lambda exonuclease in fixed ovaries, generating a stretch of single-stranded DNA (ssDNA) with 3’-OH group. A padlock probe was designed to have a unique sequence tag, which bears no homology to either the nuclear or mitochondrial genomes, flanked by 5’ and 3’ sequences complementary to the ssDNA. The two ends of the linear probe were brought to juxtaposition by hybridizing to the ssDNA, which primes rolling circle amplification after the circularization of the probe. The rolling circle amplification produced an ultralong, periodic ssDNA covalently linked to the mtDNA molecule at the restriction enzyme site (**Figure 1B**). We then performed fluorescence *in situ* hybridization targeting tag sequences to visualize the linked mtDNA therein.

*D.* melanogaster mtDNA contains a single XhoI site and a single BglII site. We designed two different padlock probes complementary to the ssDNA derived from XhoI (XhoI probe) or BglII (BglII probe) cleavage. We performed CAR assay on wild-type ovaries using either of these two probes. In ovaries, CAR signals of both probes appeared as ellipsoid-shaped puncta (**Figure 1C**). The majority of CAR signals either overlapped with or were closely associated with TFAM-mNeonGreen (**Figure 1C, 1D**), an endogenously tagged TFAM fusion protein that marks mtDNA nucleoids (Zhang *et al*, 2024). A minor fraction of CAR puncta did not colocalize with TFAM (**Figure 1C, 1D**), likely labelling uncompacted mtDNA molecules that lack TFAM coating (Isaac *et al*, 2024). Overall, 52±4% of mitochondrial nucleoids were labeled by the XhoI probe (**Figure 1E**), while 42±4% of nucleoids were labeled by the BglII probe (**Figure 1E**), demonstrating comparable efficiency of these two probes in labeling mtDNA.

To assess the specificity of the CAR assay, we applied both padlock probes to ovaries carrying either *mt:ND2^ins^* or *mt:CoI^T300I^* genome, which lacks the BglII site and the Xho site, respectively (Xu, 2008). In *mt:ND2^ins^* ovaries, the XhoI probe effectively labeled mitochondrial nucleoids, whereas CAR puncta from the BglII probe were rare, accounting for less than 2% of all puncta detected in the same ovariole (**Figure 2A, 2E**). Similarly, in *mt:CoI^T300I^* ovaries, CAR puncta from the XhoI probe were less than 2% of all puncta in the same ovariole (**Figure 2A, 2E**). These results demonstrate that CAR is both effective and specific in distinguishing mtDNA variants with SNPs *in situ*.

**Figure 2.**
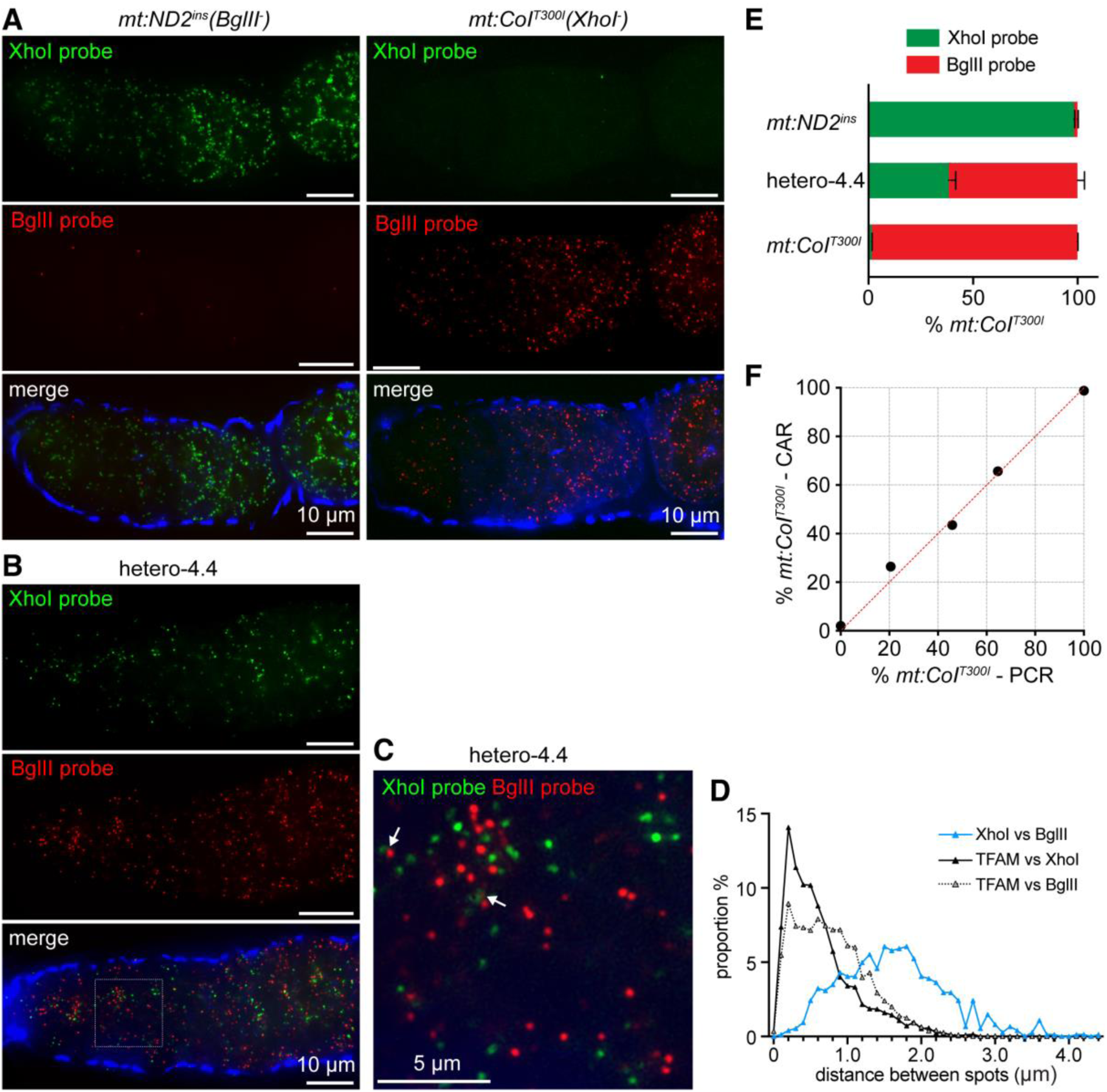
Specificity and accuracy of the CAR assay in distinguishing *mt:ND2^ins^* and *mt:CoI^T300I^*mtDNA variants *in situ*. (**A**) The CAR assay specifically detects different mtDNA variants. Ovaries from homoplasmic *mt:ND2^ins^* (left) or *mt:CoI ^T300I^* (right) flies cultured at 18°C were hybridized with both XhoI and BglII padlock probes. The XhoI probe predominantly detects *mt:ND2^ins^* , while BglII probe mainly detects *mt:CoI^T300I^*. Phalloidin (blue) stains actin. Images are projections of 10 z-stacks (0.3 μm/stack). Scale bar, 10 μm. (**B**) The CAR assay distinguishes *mt:ND2^ins^* (XhoI probe) and *mt:CoI^T300I^* (BglII probe) in heteroplasmic flies (hetero-4.4). Scale bar, 10 μm. (**C**) Magnified view of the boxed region in (**B**), showing mitochondrial nucleoids detected by both probes. White arrows indicate that signals from the two mtDNA variants are adjacent but non-overlapping. Scale bar, 5 μm. (**D**) Frequency distribution plot of distances between the two mtDNA variants in heteroplasmic flies. The shortest distances between the centers of CAR signals detected by XhoI and BglII probes were measured (n=743, blue). Bin size, 0.1 μm. (**E**) Quantification of CAR signals detected by XhoI and BglII probes in homoplasmic *mt:ND2^ins^* (n=5), heteroplasmic (hetero-4.4) flies (n=6), and homoplasmic *mt:CoI^T300I^*flies (n=5). Data are normalized to the detection efficiency of each probe and are presented as mean± SD. (**F**) CAR-based heteroplasmy quantification aligns with PCR-based measurements. Scatter plot comparing heteroplasmy levels quantified from the same fly using CAR and PCR assays. Each data point represents the mean percentage of *mt:CoI^T300I^* in each line.

### CAR accurately detects the level of heteroplasmy

To evaluate the accuracy of CAR in assaying heteroplasmy, we generated a series of heteroplasmic lines carrying both *mt:ND2^ins^* and *mt:CoI^T300I^* genomes at varying ratios using germ plasm transplantation. We performed CAR in ovaries of heteroplasmic flies cultured at 18°C using both XhoI and BglII probes, to visualize corresponding mtDNA variants simultaneously (**Figure S2A**). CAR signals of both genomes were evident, with each CAR punctum appearing spatially distinct from the other (**Figure 2B, 2C, 2D, Figure S2B**). This observation is consistent with the notion that mtDNA from different nucleoids rarely intermix (Gilkerson *et al*, 2008). We quantified the number of CAR puncta for each genome and deduced the heteroplasmy levels by factoring in the detecting efficiency of each probe (**Figure 2E**). The heteroplasmy levels determined by quantifying CAR puncta closely matched those obtained from the quantification of digested PCR products (**Figure 2F, Figure S2C, S2D**), validating the accuracy of CAR in assessing heteroplasmy levels.

### Selection against deleterious mtDNA variants takes place in differentiating cysts

The homoplasmic *mt:CoI^T300I^* flies exhibit temperature-sensitive lethality (Chen *et al*., 2015; Hill *et al*., 2014), whereas homoplasmic *mt:ND2^ins^* flies are largely healthy (Burman *et al*., 2014; Ma *et al*., 2014). To assess the mtDNA selection in heteroplasmic fly, we performed a temperature shifting assay (Hill *et al*., 2014; Ma *et al*., 2014), and quantified the load of *mt:CoI^T300I^* in progeny produced by a single heteroplasmic mother at either a restrictive temperature of 29°C or a permissive temperature of 18°C by PCR-digestion method. The load of *mt:CoI^T300I^* was lower in progeny produced at 29°C compared to their siblings produced at 18°C (**Figure 3A**). In addition, over multiple generations, the proportion of *mt:CoI^T300I^* in the population continued to decline at 29 °C, although the rate of decline reduced as *mt:CoI^T300I^* levels decreased (**Figure 3B**). These results resemble previous observations of mtDNA selection in heteroplasmic flies carrying both wild-type and *mt:CoI^T300I^* genomes (Hill *et al*., 2014; Ma *et al*., 2014), validating the use of this heteroplasmic line to investigate the mechanisms of selective inheritance.

**Figure 3.**
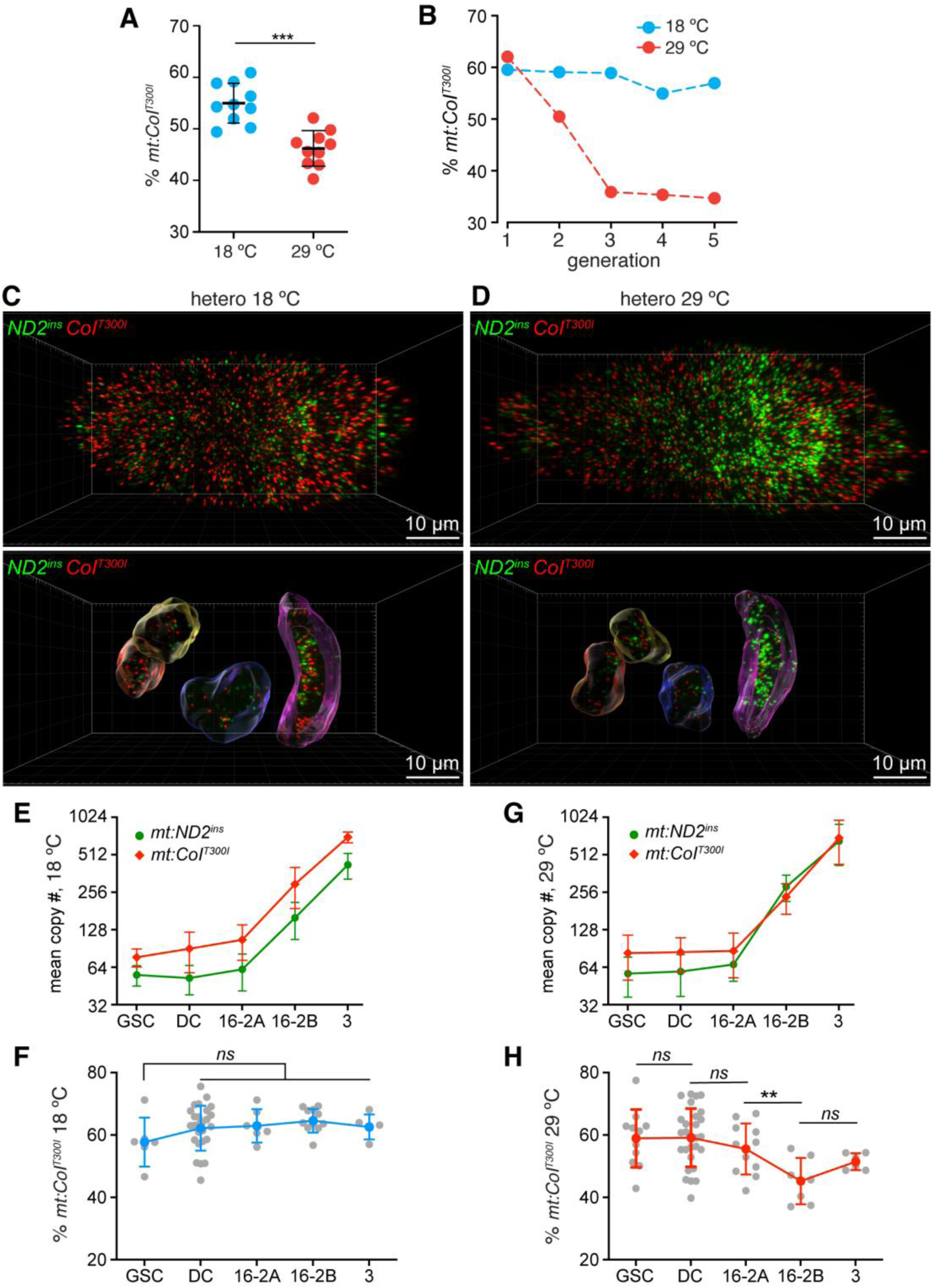
Germline selection against the *mt:CoI^T300I^* genome occurs at germarium region 2B. (**A**) Temperature shifting assay showing the heteroplasmy level of *mt:CoI^T300I^*in progeny produced at 18°C and 29°C by a single mother. Individual larvae produced at each temperature were analyzed using PCR assay (n= 10). Data are presented as mean± SD. Statistical analysis was performed using unpaired *t*-test. ***P≤ 0.001. (**B**) Proportion of *mt:CoI^T300I^* in heteroplasmic (hetero-4.4) flies maintained at 18°C or 29°C across generations. Data were quantified using PCR analysis from 20 flies per generation. (**C**) (**D**) Representative 3D rendering of CAR images from heteroplasmic flies (hetero-4.4). Flies were maintained at 18°C (**C**) or 29°C (**D**) for 4 days before dissection. The *mt:ND2^ins^* (green) and *mt:CoI^T300I^* (red) mtDNA are detected by XhoI and BglII probes, respectively. In bottom panels, 3D rendering of developing cysts and CAR signals of *mt:ND2^ins^* (green) and *mt:CoI^T300I^* (red) mtDNA within the cysts are shown. The 2-cell cyst (orange), 4-cell cyst (yellow), region 2A (blue) and region 2B (magenta) are segmented using Imaris. Scale bar, 10 μm. (**E**) (**G**) Quantification of *mt:ND2^ins^* and *mt:CoI^T300I^* mitochondrial DNA numbers at different germarium stages using the CAR assay in heteroplasmic flies maintained at 18°C (**E**) and 29°C (**G**). For each temperature condition (18°C (**E**) and 29°C (**G**)), the number of each mtDNA variant in germline stem cells (GSC, n=6, 11), dividing cysts (DC, n=25, 28), region 2A (16-2A, n=6, 11), region 2B (16-2B, n=11, 7) and region 3 (3, n=5, 6) are shown. Data are presented as mean± SD. (**F**) (**H**) Heteroplasmy levels of the of *mt:CoI^T300I^* genome at different germarium stages in heteroplasmic flies maintained at 18°C (**F**) and 29°C (**H**). The proportion of *mt:CoI^T300I^*relative to total mtDNA was quantified using the CAR assay. Data are presented as mean± SD. Statistical analysis was performed using an unpaired *t*-test. ns, not significant; **P≤ 0.01.

To monitor the dynamics of two mtDNA variants in developing germ cells, we performed CAR in ovaries of heteroplasmic flies cultured at either 18°C or 29°C (**Figure S3A, S3B**). Ovaries were co-stained with fluorescently labeled phalloidin, which binds to actin filaments and thereby outlines germ cells (Merkle, 2023), to identify development stages. We segmented germline cysts at different development stages in reconstructed 3D volumes (**Figure 3C, 3D; Movie 1, 2**) and quantified the number of CAR puncta derived from each genome.

At 18°C, the total number of nucleoids, including both genomes, was estimated to be approximately 130 in a GSC and 140 in a dividing cyst (2-, 4-, or 8-cell cyst) after factoring in the detection efficiency of each probe (**Figure 3E**). For 16-cell cysts, the copy number increased from ∼160 in region 2A to ∼443 in region 2B (**Figure 3E**). In region 3, the copy number reached ∼1134 per budding egg chamber (**Figure 3E**). This data aligns with the previous finding that mtDNA replication is largely quiescent in dividing cysts but activated in differentiating cysts at region 2B (Hill *et al*., 2014). We also quantified the load of *mt:CoI^T300I^* in each cyst or GSC, and found that proportion of *mt:CoI^T300I^* were relatively stable over all developmental stages in a germarium (**Figure 3F**), indicating a lack of selection at the permissive temperature of 18°C.

At 29°C, the total number of nucleoids was comparable to that at the same developmental stage at 18°C and remained stable in GSCs, dividing cysts, and 16-cell cysts in region 2A (**Figure 3G**). A significant increase in nucleoid numbers was also observed in the 16-cell cysts in region 2B (**Figure 3D, 3G**). However, *mt:ND2^ins^* increased to a greater extent than *mt:CoI^T300I^*, resulting a 15% reduction in *mt:CoI^T300I^* load in region 2B, compared to earlier germarium stages (**Figure 3G, 3H**). This value closely matched the 12% reduction of *mt:CoI^T300I^* observed over a single generation in population studies (**Figure 3B**). Additionally, the heteroplasmy level of *mt:CoI^T300I^* in budding egg chambers (region 3) was comparable to that in region 2B (**Figure 3H**). Together, these findings indicate the cross-generational selection against this deleterious mtDNA variant mainly occurs in the differentiating cysts at germarium region 2B.

### The healthy mitochondrial genome is preferentially replicated over the deleterious variant

The reduced mutation load could be caused by a selective elimination of mutant genome through mitophagy or other quality control mechanisms. However, the absolute number of the *mt:CoI^T300I^* genome in heteroplasmic ovaries was not reduced at any stages of developing germ cells in the germarium (**Figure 3G**), arguing against a model of active elimination. We next asked whether the faster increase of the healthy genome, *mt:ND2^ins^* inregion 2B reflects different replication frequencies between these two genomes. We combined CAR with EdU incorporation to assess the replication of *mt:ND2^ins^* and *mt:CoI^T300I^* in heteroplasmic ovaries. After a 2-hour EdU incubation, a proportion of mtDNA was labeled with EdU **(Figure 4A, 4B, Figure S4)**, consistent with previous studies showing that mtDNA undergoes relaxed replication (Birky, 1994; Bogenhagen & Clayton, 1978). Most CAR signals partially overlapped with EdU signals, likely because CAR products extend away from mtDNA loci (**Figure 1**). The proportion of replicating events, indicated by EdU-positive CAR signals, was similar between two genomes in both region 2B and budding egg chambers of heteroplasmic flies cultured at 18°C (**Figure 4A, 4C, Figure S4**). However, at 29°C, *mt:ND2^ins^* exhibited a markedly higher replication frequency than *mt:CoI^T300I^* in region 2B, but not in region 3 (**Figure 4B, 4D, Figure S4**). Together, these results provide direct evidence that the healthy mitochondrial genome in fact replicates at a higher frequency than the co-existing deleterious variant.

**Figure 4.**
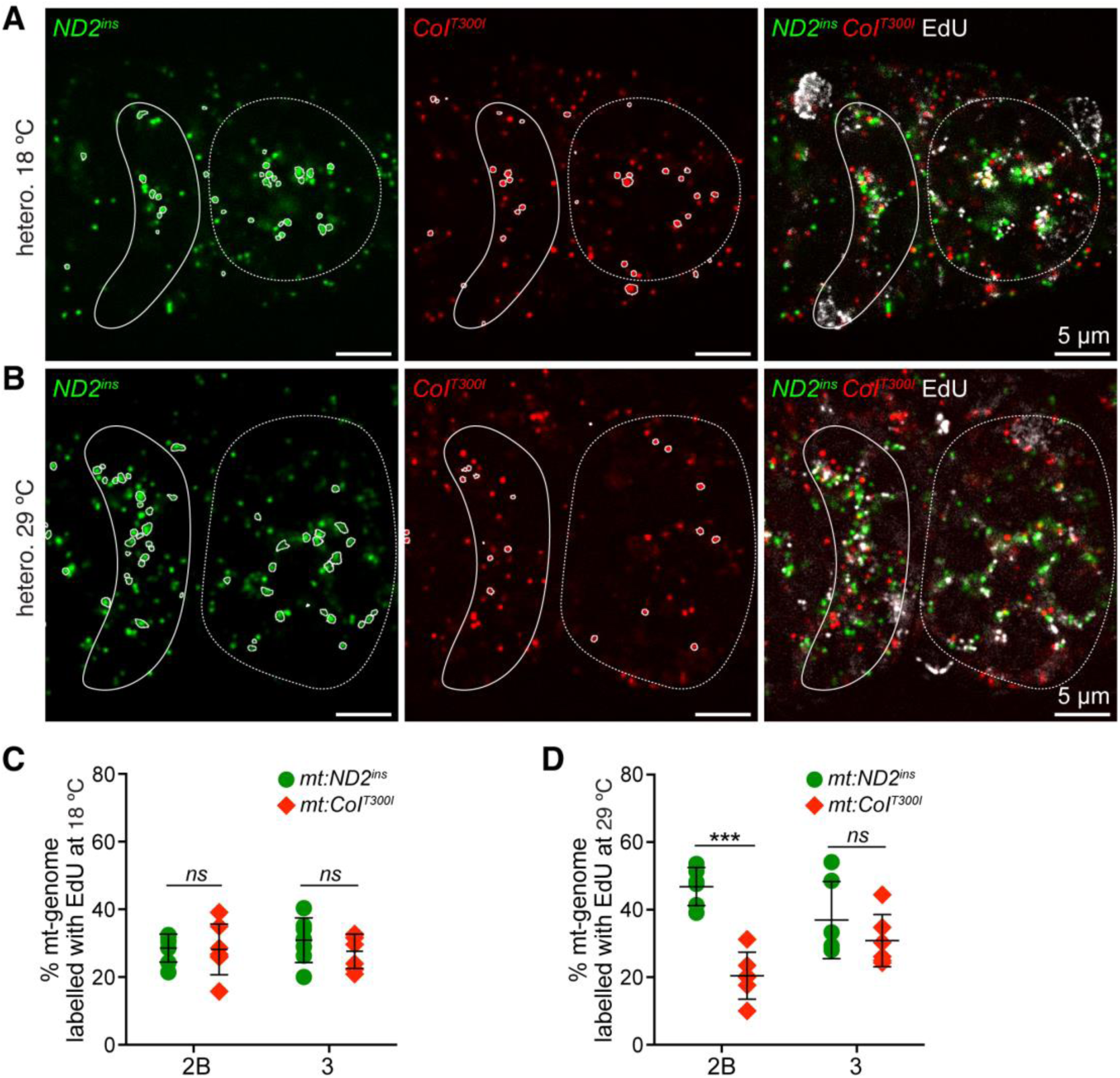
The *mt:ND2^ins^*genome replicates more efficiently than the deleterious *mt:CoI^T300I^* genome at germarium stage 2B. (**A**) (**B**) Representative images of simultaneous EdU staining and CAR assay in heteroplasmic ovaries. The flies were maintained at 18°C (**A**) or 29°C (**B**) for 4 days before dissection. The *mt:ND2^ins^* (green) and *mt:CoI^T300I^* (red) mtDNA are detected by XhoI and BglII probes, respectively. Replicating mtDNA is labeled with EdU (white). CAR signals enclosed by white lines indicate those colocalizing with EdU staining. Germarium region 2B (solid line) and region 3 (dashed line) are indicated. Scale bar, 5 μm. (**C**) (**D**) Quantification of replicating mtDNA for each mtDNA variant in heteroplasmic flies maintained at 18°C (**C**) and 29°C (**D**). The proportion of replicating *mt:ND2^ins^* and *mt:CoI^T300I^* genomes was determined by normalizing EdU-positive CAR signals to the total number of corresponding mtDNA signals under each temperature in germarium region 2B (n=7, 6) and budding egg chambers (region 3, n=7, 6). Statistical analysis was performed using an unpaired *t-*test. ns, not significant; ***P≤ 0.001.

## DISCUSSION

A padlock probe-mediated rolling-circle amplification method was previously developed to detect mtDNA SNPs in cultured cells (Larsson *et al*., 2004). In this approach, wild-type and mutant padlock probes compete for the same target sequence, and the long single-stranded (ssDNA) 3’ overhang generated after exonuclease treatment could potentially form secondary structures, further limiting its efficiency in detecting mtDNA SNPs in tissues. In *Drosophila*, most mtDNA mutations affect certain restriction enzyme cleavage sites. We implement the CAR assay by leveraging this feature to generate distinct, short ssDNA stretches, derived from different enzyme cleavage sites on each genome. Compared to the 10% efficiency observed in cultured cells using the previous method (Larsson *et al*., 2004), CAR displayed 4-5 fold increase in detection efficiency (**Figure 1E**). In addition, the rolling cycle amplification in CAR is initiated at the unique sequences on each genome, minimizing non-specific amplification and thereby further enhancing the specificity. We also found that treating ovaries with Triton X-100, rather than ethanol or proteases, was sufficient for tissue permeabilization while better preserving tissue integrity and making CAR compatible with immunofluorescence antibody staining (**Methods**). Another approach integrating CRISPR-associated (Cas) proteins with RCA amplification requires complex DNA/RNA probes (Zhang *et al*., 2018), and were ineffective in spatially distinguishing different mtDNA genotypes in *Drosophila* ovaries (**Figure S1**). Notably, CAR products are covalently linked to endogenous mtDNA, enabling single-molecule resolution of heteroplasmy within tissues. Most human disease-associated mtDNA mutations are point mutations (Taylor & Turnbull, 2005). While CAR assay targets SNPs at restriction enzyme sites, recent advances in CRISPR/Cas, ZFN and TALEN based DNA editing technologies hold the potential to expand this approach to visualize any point mutations on mtDNA or the nuclear genome. Hence, CAR would offer unprecedented spatial resolution to track mtDNA segregation and dynamics in tissues, which is crucial for understanding the pathogenic mechanisms of mtDNA diseases.

Using CAR, we demonstrated that the healthier genome *mt:ND2^ins^* exhibited a higher frequency of replication than *mt:CoI^T300I^* within the same differentiating germline cyst. This data provides direct evidence that healthy genome has an advantage in replication at this stage and thereby outcompetes the deleterious variants. The reduction of *mt:CoI^T300I^* levels in germarium region 2B was comparable to its decline across a single generation, indicating that the selection primarily occurs at this stage. Consistent with this notion, the frequency of replication was comparable between *mt:ND2^ins^* and *mt:CoI^T300I^* in budding egg chambers, and no further reduction of *mt:CoI^T300I^* was observed beyond region 2B. In female germarium, mtDNA copy number is ∼100 in a GSC and remains stable in dividing cysts (**Figure 3E, 3G**). From region 2B onward, germ cells undergo 16.6 replication cycles, ultimately accumulating about 10 million copies of mtDNA in a mature oocyte (Chen *et al*, 2025; Wolff *et al*, 2013). Our data indicates that the healthier genome, *mt:ND2^ins^* underwent ∼2.1 replication cycles, compared to ∼1.4 cycles for *mt:CoI^T300I^* in region 2B. This modest difference in replication frequency sufficiently accounts for the ∼15% reduction of *mt:CoI^T300I^* that is commonly observed across one generation in heteroplasmic flies. This reasoning not only corroborates with the finding that replication competition occurs in a narrow developmental window but also underscores the robustness of the replication competition in limiting the transmission of deleterious mtDNA mutations.

The absolute number of *mt:CoI^T300I^* was not reduced throughout germ cell development under the restrictive condition (**Figure 3G**), arguing against the idea that the alteration in mtDNA heteroplasmy during cyst differentiation is directly driven by the selective removal of deleterious mtDNA variants *via* mitophagy or other quality control processes. Consistent with this notion, programed germline mitophagy in dividing cysts, which promotes mtDNA selective inheritance, occurs regardless of the presence mtDNA mutations (Palozzi *et al*., 2022). Noteworthy, knocking down Atg1 or BNIP3, essential factors in PGM, reduces wild-type mtDNA levels while compromising the selective inheritance (Palozzi *et al*., 2022). It would be interesting to test whether PGM might regulate mtDNA replication to indirectly influence mtDNA selection.

Prior to region 2B, mitochondrial fission allows effective mtDNA segregation to minimize complementation between wild-type and deleterious mtDNA alleles (Chen *et al*., 2020; Lieber *et al*., 2019), setting the stage for selection based on the functionality of individual genomes. At region 2B, mtDNA expression serves as a stress test for each genome’s integrity—wild-type genomes would produce functional respiratory chain complexes (RC), whereas deleterious mtDNA variants would lead to defective RC. In ovaries, many nuclear-encoded mitochondrial proteins, including key mtDNA replication factors, are synthesized on the mitochondrial outer membrane. This local translation, along with the import of preproteins into the mitochondrial matrix relies on the mitochondrial membrane potential generated by mitochondrial respiration. Therefore, in heteroplasmic germ cells, mitochondria carrying the functional genome would have an advantage over those containing deleterious mutations in acquiring replication factors. As a result, wild-type or healthier genomes are more likely to replicate and outcompete coexisting defective counterparts. Consistent with this model, partial reduction of replication factors such as PolG1has been shown to intensify this competition, thereby enhancing the selective inheritance of functional mtDNA (Chiang *et al*, 2019; Palozzi *et al*, 2022). Conversely, when mtDNA replication is severely impaired across all mitochondria, the efficiency of purifying selection is diminished mtDNA selection (Zhang et al, 2019). All these observations underscore the critical link between mtDNA replication and selective inheritance. In *C. elegans*, an mtDNA variant carrying a large deletion gains a selfish replicative advantage by preferentially associating with the mtDNA replicase, facilitating its stable transmission across generations (Yang *et al*., 2022). It would be interesting to test whether any of the mtDNA replication factors are more enriched in healthy organelles, which might be the molecular underpins of replication competition and selective inheritance.

Purifying mtDNA selection, potentially at the organelle or genome level, has been observed during primordial germ cell migration in humans (Floros *et al*., 2018) and folliculogenesis in mice models (Ru *et al*., 2024). In both cases, selection against deleterious mtDNA mutations coincided with the upregulation of genes involved in mtDNA replication, transcription (Floros *et al*., 2018) or mitochondrial protein translation (Ru *et al*., 2024). Therefore, replication competition might represent a conserved mechanism that enforce the selective mtDNA inheritance and shape the evolution of mitochondrial genome in metazoan.

## SUPPLEMENTARY FIGURES AND FIGURE LEGENDS

**Supplementary Figure 1.**
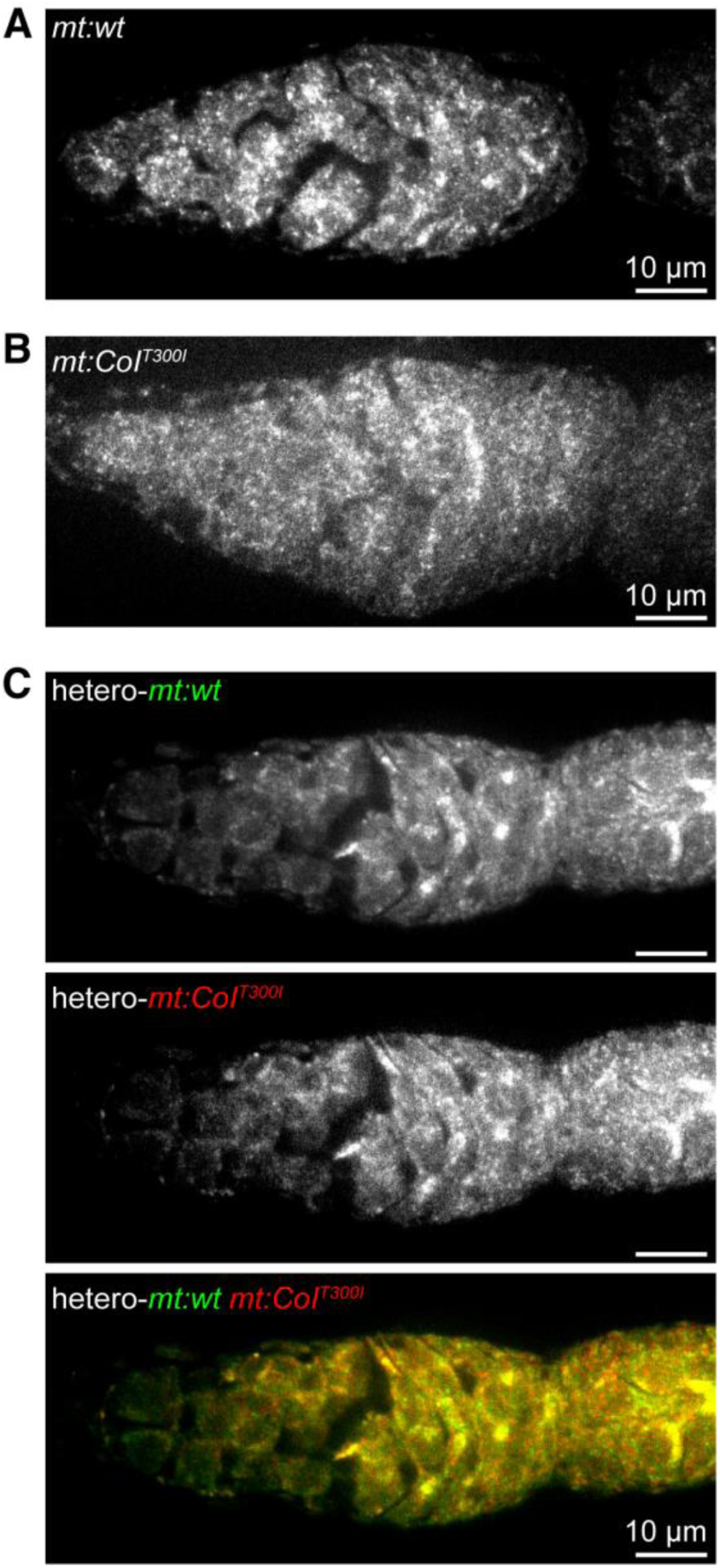
The CasPLA method cannot detect mtDNA variants in *Drosophila* ovaries. (**A**)-(**C**) Representative images of the CasPLA method (Zhang *et al*., 2018) applied to detect mtDNA in wild-type (**A**), homoplasmic *mt:CoI^T300I^* (**B**), and heteroplasmic (**C**) flies. Fluorophore-labeled ssDNA probes were designed to detect *mt:wt* (green) and *mt:CoI^T300I^* (red) genome. Scale bar, 10 μm.

**Supplementary Figure 2.**
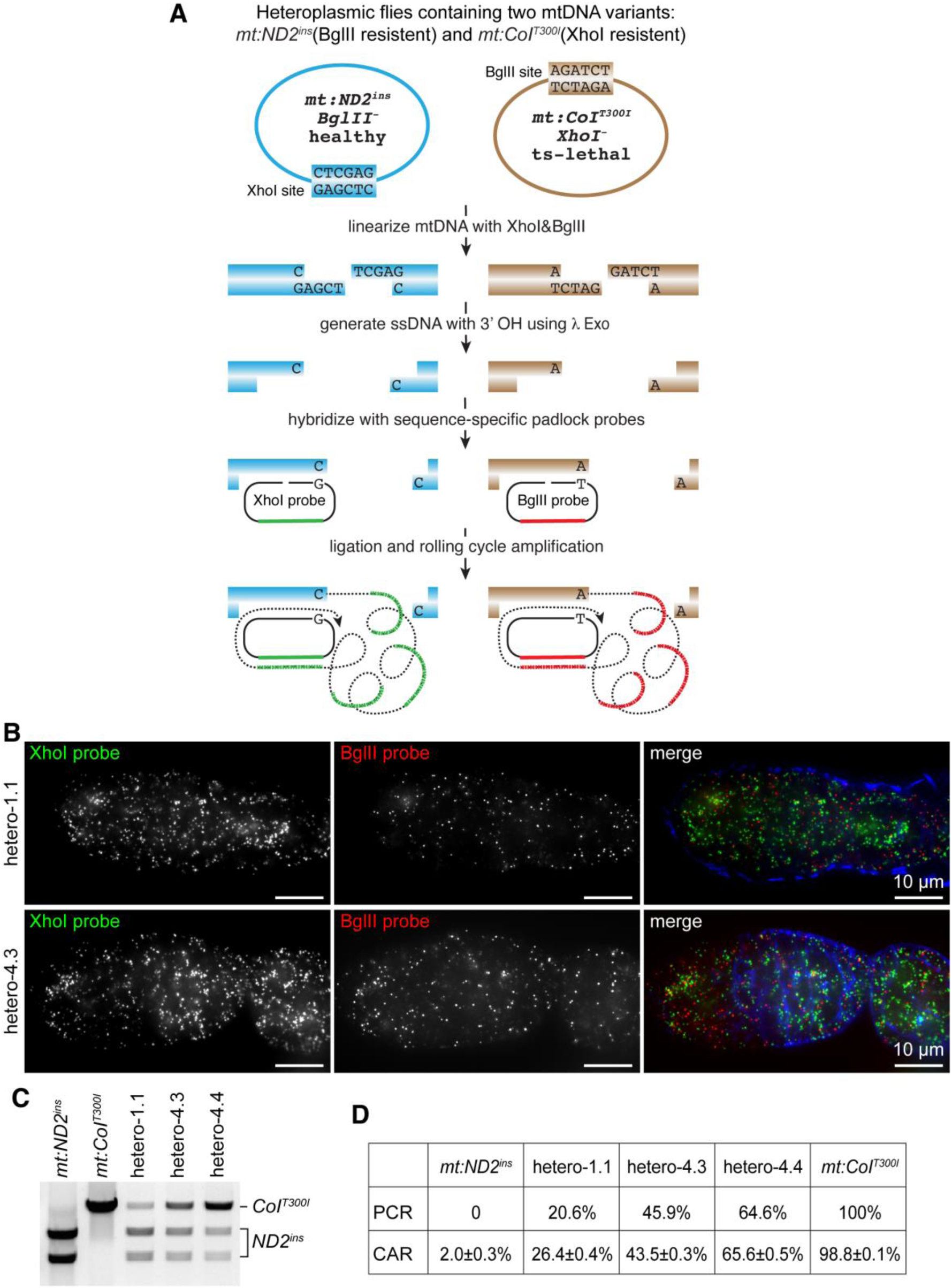
The heteroplasmy levels quantified using CAR assay are consistent with those derived from PCR assay. **(A)** Scheme of CAR assay for distinguishing mtDNA variants in the heteroplasmic fly. The mitochondrial genome variants, *mt:ND2^ins^* (blue) and *mt:CoI^T300I^* (brown), abolish the BglII and XhoI sites on mtDNA, respectively. Homoplasmic *mt:ND2^ins^* flies are as healthy as wild-type flies (Burman *et al*., 2014; Ma *et al*., 2014), whereas homoplasmic *mt:CoI^T300I^* flies exhibit temperature sensitive lethality (ts-lethal) (Chen *et al*., 2015; Hill *et al*., 2014). In heteroplasmic flies carrying these two mtDNA variants, endogenous mtDNA is digested by XhoI and BglII restriction enzymes, followed by Lambda exonuclease (λ Exo) treatment to generate single-stranded DNA (ssDNA). The XhoI and BglII padlock probes are designed with a DNA sequence complementary to the ssDNA adjacent to XhoI and BglII restriction site, respectively. After annealing, circular ligation, and rolling circle amplification, the CAR products are visualized by fluorescence *in situ* hybridization using probes specific to the unique sequence tags on each padlock probe. (**B**) Representative CAR images of the germarium from heteroplasmic flies (hetero-1.1 and hetero-4.3). Flies were maintained at 18°C for 4 days before dissection. The *mt:ND2^ins^* (green) and *mt:CoI^T300I^*(red) mtDNA are detected by XhoI and BglII probes, respectively. Phalloidin (blue) stains actin. Images are projections of 10 z-stacks (0.3 μm/stack). Scale bar, 10 μm. (**C**) Quantification of heteroplasmy levels using the PCR method. A 4-kb PCR fragment spanning the *mt:CoI* site was amplified from total DNA extracted from ten flies of indicated genotypes. A DNA gel of PCR fragments following XhoI enzyme digestion is shown. The *mt:ND2^ins^* genome is cleaved by XhoI, producing two smaller bands, whereas *mt:CoI^T300I^* is resistant to XhoI digestion. Homoplasmic *mt:ND2^ins^*, homoplasmic *mt:CoI^T300I^* and heteroplasmic flies (hetero-1.1, hetero-4.3, and hetero-4.4) were analyzed. (**D**) Quantification of heteroplasmy levels using the CAR assay in 2B cysts is consistent with those obtained using the PCR method. The data represent mean± SD.

**Supplementary Figure 3.**
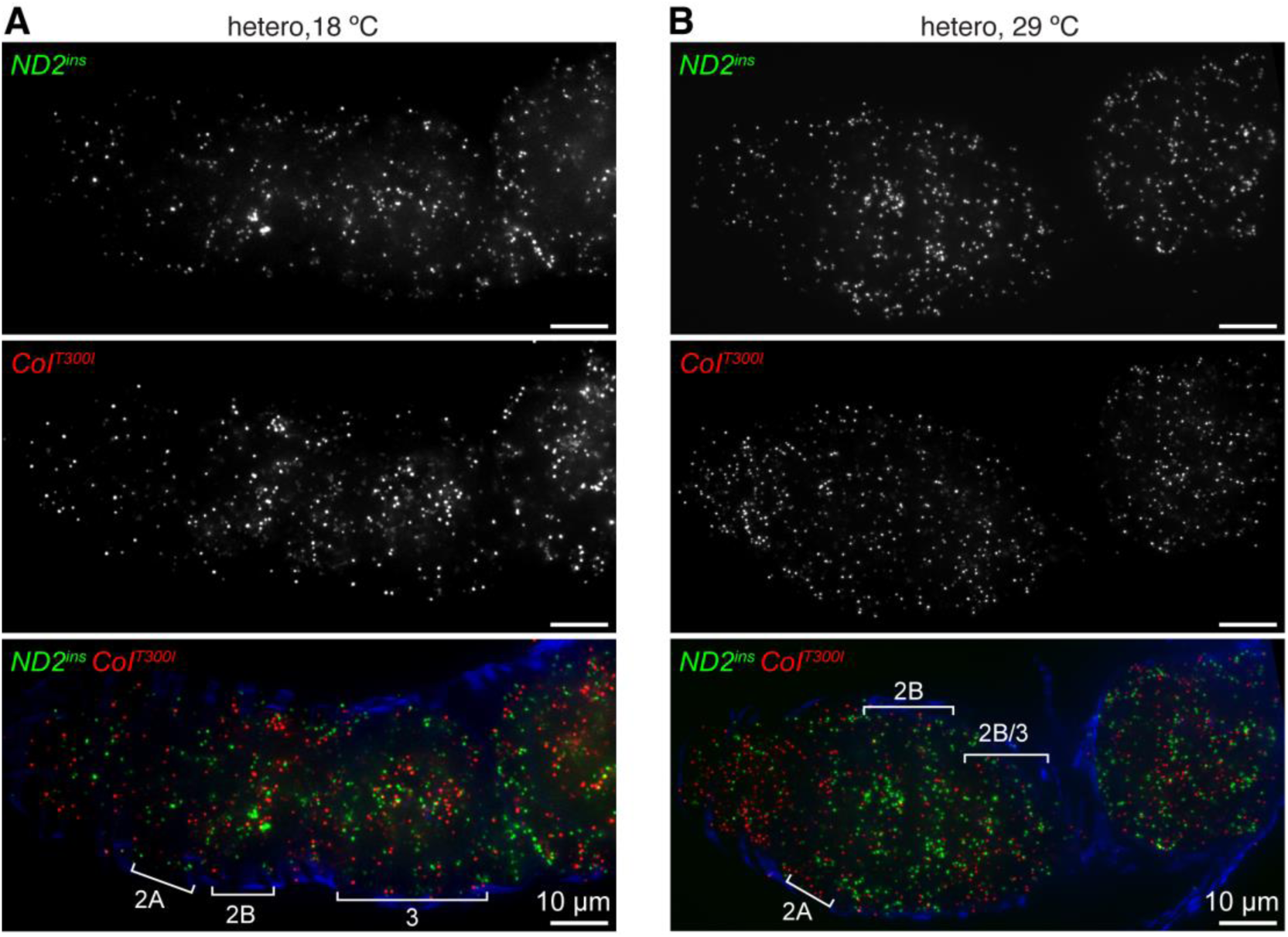
Representative CAR images of the germarium from heteroplasmic (hetero-4.4 flies) flies. Flies were maintained at 18°C (**A**) or 29°C (**B**) for 4 days before dissection. Germarium regions 2A and 2B of 16-cell cysts, as well as budding egg chambers (region 3), are indicated. The *mt:ND2^ins^* (green) and *mt:CoI^T300I^*(red) mtDNA are detected by XhoI and BglII probes, respectively. Phalloidin (blue) stains actin. Images are projections of 10 z-stacks (0.3 μm/stack). Scale bar, 10 μm.

**Supplementary Figure 4.**
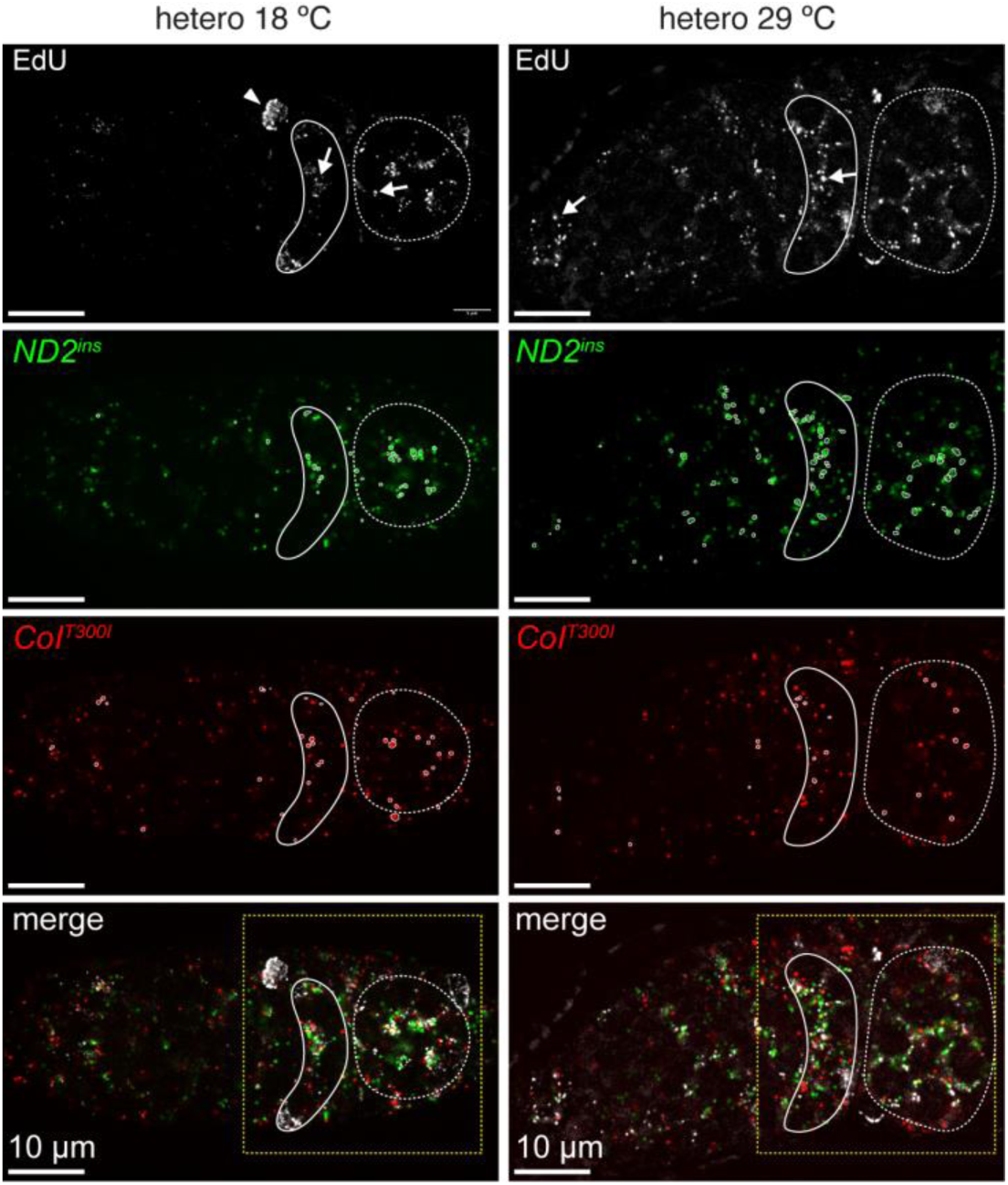
The *mt:ND2^ins^*genome replicates more efficiently than the deleterious *mt:CoI^T300I^*genome at germarium stage 2B. Representative images of simultaneous EdU staining and CAR assay in heteroplasmic germarium. Flies were maintained at 18°C or 29°C for 4 days before dissection. The *mt:ND2^ins^* (green) and *mt:CoI^T300I^* (red) mtDNA are detected by XhoI and BglII probes, respectively. Replicating mtDNA is labeled with EdU (white). CAR signals circled by white lines are those colocalized with EdU staining. Germarium region 2B (solid line) and region 3 (dashed line) are outlined. The arrowhead points to a replicating nuclear DNA and arrows indicate replicating mtDNA. The boxed region in the merged panels is magnified and shown in Figure 4. Scale bar, 10 μm.

## MOVIE LEGENDS

**Movie 1. 3D rendering of CAR images and germarium cysts from heteroplasmic flies maintained at 18°C.** The *mt:ND2^ins^* (green) and *mt:CoI^T300I^* (red) mtDNA are detected by XhoI and BglII probes, respectively. The 2-cell cyst (orange), 4-cell cyst (yellow), region 2A (blue) and region 2B (magenta) are shown. Scale bar, 5 μm.

**Movie 2. 3D rendering of CAR images and germarium cysts from heteroplasmic flies maintained at 29°C.** The *mt:ND2^ins^* (green) and *mt:CoI^T300I^* (red) mtDNA are detected by XhoI and BglII probes, respectively. The 2-cell cyst (orange), 4-cell cyst (yellow), region 2A (blue) and region 2B (magenta) are shown. Scale bar, 5 μm.

## MATERIALS AND METHODS

### Fly stocks and husbandry

*Drosophila melanogaster* stocks were reared in a humidity-controlled incubator under a 12-hour light-dark cycle using standard cornmeal molasses agar media. Heteroplasmic *Drosophila melanogaster* strains were maintained at 18 °C unless otherwise indicated. The following fly stocks reared at 25 °C were used: TFAM-mNeonGreen (Zhang *et al*., 2024), *w^1118^* (*mt*:*CoI^T300I^*) (Xu, 2008), *w^1118^* (*mt*:*ND2^ins^*) (Xu, 2008).

### Generation of heteroplasmic flies

Heteroplasmic flies were generated using germplasm transplantation, as previously described (Hill *et al*., 2014; Ma *et al*., 2014). Briefly, germplasm from homoplasmic mt:*ND2^ins^* embryos were injected into the posterior end of homoplasmic *mt:CoI^T300I^* recipient embryos. Hatched females from injected embryos were crossed to *w^1118^* males, and the progeny from this cross were subsequently mated *en masse* to *w^1118^* males at room temperature. To select for heteroplasmic flies, embryos were shifted to 29°C, and surviving progeny were identified as heteroplasmic flies. Female escapers were individually crossed with *w^1118^* males and maintained as founder lines. The proportion of *mt:CoI^T300I^* in each founder line was quantified (see below) . Three lines with low (hetero-1.1), medium (hetero-4.3), and high (hetero-4.4) levels of *mt:CoI^T300I^* were chosen and maintained at 18°C.

### Quantification of heteroplasmy using PCR

Total DNA was extracted from the indicated flies using the DNeasy Blood & Tissue kit (QIAGEN). The BglII site is located 1568 base pairs (bp) upstream of the XhoI site on mtDNA. A 4 kb mtDNA fragment spanning both XhoI and BglII sites was amplified from total DNA using PCR primers described previously (Hill *et al*., 2014). The PCR products were digested overnight with the XhoI restriction enzyme and separated by agarose gel electrophoresis. DNA band intensities were quantified using ImageJ software. The heteroplasmy level was determined as the proportion of XhoI-resistant DNA fragment (mt:*CoI^T300I^*) relative to the total DNA.

### mtDNA selection assay

The mtDNA selection assay was performed following previously established methods (Hill *et al*., 2014). To examine mtDNA selection across generations, progeny from the hetero-4.4 founder line were divided into two groups and maintained at 18°C and 29°C, respectively. In each generation, 50 flies (mixed male and female) were randomly selected for mating. After 5 days, these flies were collected and stored at −80°C until heteroplasmy levels were quantified. Flies were maintained for at least five generations at respective temperature.

To assess mtDNA selection during female germline development, a single heteroplasmic female was mated with multiple *w^1118^* males at 18°C for 3 days. The flies were then transferred to a new vial and acclimated at 29°C for 4 days. During this period, embryos were discarded to ensure that only germ cells that developed at 29°C were analyzed. Progeny produced on days 1-3 at 18°C and days 5–7 at 29°C were collected, and their heteroplasmy levels were quantified.

### CAR (Circular DNA Amplification at Restriction enzyme sites) assay in *Drosophila* ovaries

Flies with specified genotypes were maintained at 18°C, 25°C, or 29°C, depending on experiment requirements. Ovaries were dissected in Schneider’s *Drosophila* medium supplemented with 10% fetal bovine serum (FBS) and fixed in 4% paraformaldehyde for 20 minutes. Following fixation, tissues were washed three times in 1×PBST (PBS with 0.1% Tween-20) and permeabilized in RIPA buffer (150 mM NaCl, 1% TritonX-100, 0.5% sodium deoxycholate, 0.1% SDS, 1 mM EDTA, 50 mM Tris-HCl pH 8.0) for 30 minutes at room temperature with gentle rotation. The permeabilization step was repeated three times using fresh RIPA buffer, followed by two washes in 1×PBST. To prevent over-treatment by RIPA, tissues were fixed again in 4% paraformaldehyde for 30 min, followed by three additional washes in 1×PBST.

For restriction enzyme digestion, ovaries were incubated with indicated NEB restriction enzyme (XhoI, BglII or both) in NEBuffer r3.1 at 37°C overnight. The next day, the restriction enzyme was removed by three washes in 1×PBST, and tissues were treated with Lambda exonuclease for 15 minutes at 37°C, followed by four additional washes in 1×PBST. Padlock probe hybridization was then performed by incubating tissues overnight at 16°C in the probe-ligase mixture (2 μM Padlock probe, 20 units/μl T4 DNA ligase, 1× T4 DNA ligase buffer). The next day, samples were washed twice in 1×PBST (pre-cold on ice) for 3 minutes each before proceeding to rolling cycle amplification (RCA). For RCA, tissues were incubated in 250 μl EquiPhi29 reaction mix (0.25 U/μl EquiPhi29, 1 mM DTT, 0.5 mM dNTPs, 1× EquiPhi29 reaction buffer) at 30°C for 4-6 hours with gentle rotation, followed by washing three times in PBS for 5 minutes each.

To visualize the amplified signals, fluorescent probe *in situ* hybridization was performed. Tissues were first pre-incubated in hybridization buffer (10% deionized formamide, 2×SSC, 10% dextran sulfate) at 37°C for 15 minutes, followed by incubation in hybridization mix (10% deionized formamide, 2×SSC, 10% dextran sulfate, 2.5 μl salmon sperm DNA, 200 nM fluorophore-labeled detection probes) at 37°C for at least 7 hours or overnight in the dark. The next day, tissues were sequentially washed at 37°C in 2×SSC buffer (10% deionized formamide, 2×SSC), 1×SSC buffer (10% deionized formamide, 1×SSC) and 0.5×SSC buffer (10% deionized formamide, 0.5×SSC) for 15 minutes each, before being immersed in PBS for 5 minutes. For phalloidin staining, tissues were incubated with CF®405M Phalloidin for 1 hour at room temperature, followed by three washes in PBS for 7 minutes each. Finally, ovaries were mounted using VECTASHIELD antifade mounting medium and imaged using a Nikon CSU-W1 SoRa confocal system (Nikon SR Plan Apo IR 60×/1.27 oil lens; Nikon Element software; Yokogawa CSU-W1 SoRa Confocal Scanner Unit). Image deconvolution was performed using NIS Elements AR (6.02.03) software for further processing.

### Simultaneous EdU staining and CAR assay

Newly eclosed female flies with the specified genotypes were reared for 5 days at the indicated temperature. Ovaries were dissected in Schneider’s *Drosophila* medium supplemented with 10% FBS and transferred to fresh medium containing 7 μM aphidicolin, followed by incubation for 3 hours at the respective temperature. The medium was then replaced with fresh medium containing 20 μM EdU and 7 μM aphidicolin, and incubation continued for an additional 2 hours at 18°C or 29°C. After two 5-minute washes in *Drosophila* medium, ovaries were fixed in 4% paraformaldehyde in PBS for 20 minutes at room temperature. The subsequent procedures, including permeabilization, second fixation, restriction enzyme digestion, and Lambda exonuclease treatment, were performed as described in the CAR assay protocol.

The Padlock probe ligation step was adapted for EdU staining by incubating tissues in probe-ligase mixture (2 μM Padlock probe, 20 units/μl HI-T4 DNA ligase, 1×HI-T4 DNA ligase buffer) for 3 hours at 25°C, followed with two 3-minute washes with 1×PBST. After rolling circle amplification, tissues were immersed in 3% BSA in 1×PBS for 5 minutes. EdU detection was performed according to the manufacturer’s instructions using the Click-iT EdU labeling kit (Alexa Fluor 488 dye). Following the Click-iT reaction, tissues were rinsed once with 3% BSA in PBS and then washed overnight in 3% BSA in PBS in the dark.

The subsequent steps followed the fluorescent probe *in situ* hybridization protocol of the CAR assay. Then ovaries were mounted on slides using VECTASHIELD antifade mounting medium with DAPI, and images were acquired using a Nikon CSU-W1 SoRa confocal system. Image deconvolution was performed using NIS Elements AR (6.02.03) software for further processing.

### Quantification of mtDNA variants in the germarium using CAR assay

To quantify the absolute numbers of two mtDNA variants at different developmental stages, z-stack images (0.3 μm/section) from the CAR assay were processed using Imaris (version 9.7.0). Mitochondrial nucleoids labeled with BglII and XhoI probes were detected using the “Spot” function in the “Cy3” and “Alexa488” channels, respectively. For each channel, spots were created using the “automatic creation” algorithm with the “different spot sizes (region growing)” option. The estimated XY and Z diameters were set to 0.4 μm and 0.8 μm, respectively. The “background subtraction” method was applied for spot detection, and spots with a quality score lower than 15 were filtered out.

To outline individual cysts, the “Surface” function was used. In the phalloidin channel, a surface was created using the “Add Contour” function, and cysts at different developmental stages were identified in 3D based on cell size and the number of ring canals stained with phalloidin. Once contours were drawn, the “Create Surface” function was applied to generate a surface for each cyst. The number of BglII or XhoI-labeled spots within a cyst was quantified using the “Shortest Distance to Surface” filter, with spots having a distance less than 0 considered inside the cyst. The number of each mtDNA variant was then determined by normalizing spot counts to the detection efficiency of their respective probes.

To quantify the distances between TFAM and BglII, TFAM and XhoI or BglII and XhoI-labeled mitochondrial nucleoids in the germarium, spots were generated for each channel using the method described above. The “Shortest Distance to Spots” statistics were obtained under the “Statistics” function for each spot pair and used for calculation and plotting.

### Quantification of EdU-positive CAR signals

EdU-positive CAR signals in the germarium were quantified using ImageJ (v2.16.0). Images were pre-processed using the “Deconvolution” and “Subtracting background” functions, then split into single channel images. The DAPI channel was used to delineate different developmental stages. For the BglII or XhoI probe channels, individual puncta were segmented using the “Adjust Threshold” function. Connected objects were separated using the “Watershed” and “Brush” tools. The “Analyze Particles” tool was then used to identify and analyze objects, with those smaller than 8.5 pixels filtered out. Next, segmented objects were superimposed onto the EdU channel, and objects without any overlap with the EdU signal were removed using the “ROI manager”. The ratio of EdU-positive CAR signals was calculated by determining the proportion of CAR signals that were either completely or partially overlapped with EdU relative to the total number of CAR signals, in the indicated germarium stages.

### Statistical analyses

Two-tailed Student’s t-test was used for statistical analyses. The difference was considered statistically significant when p<0.05. Results are represented as mean±SD of the number of determinations.

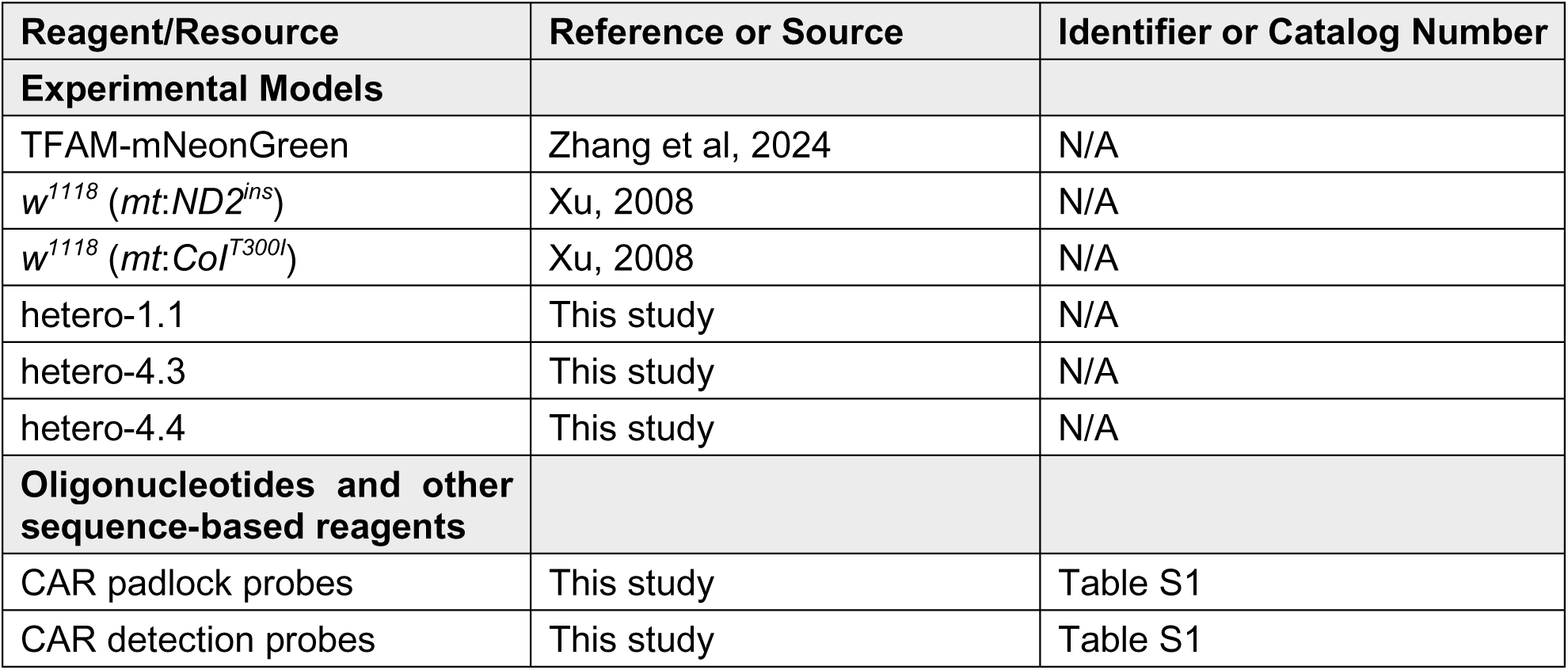
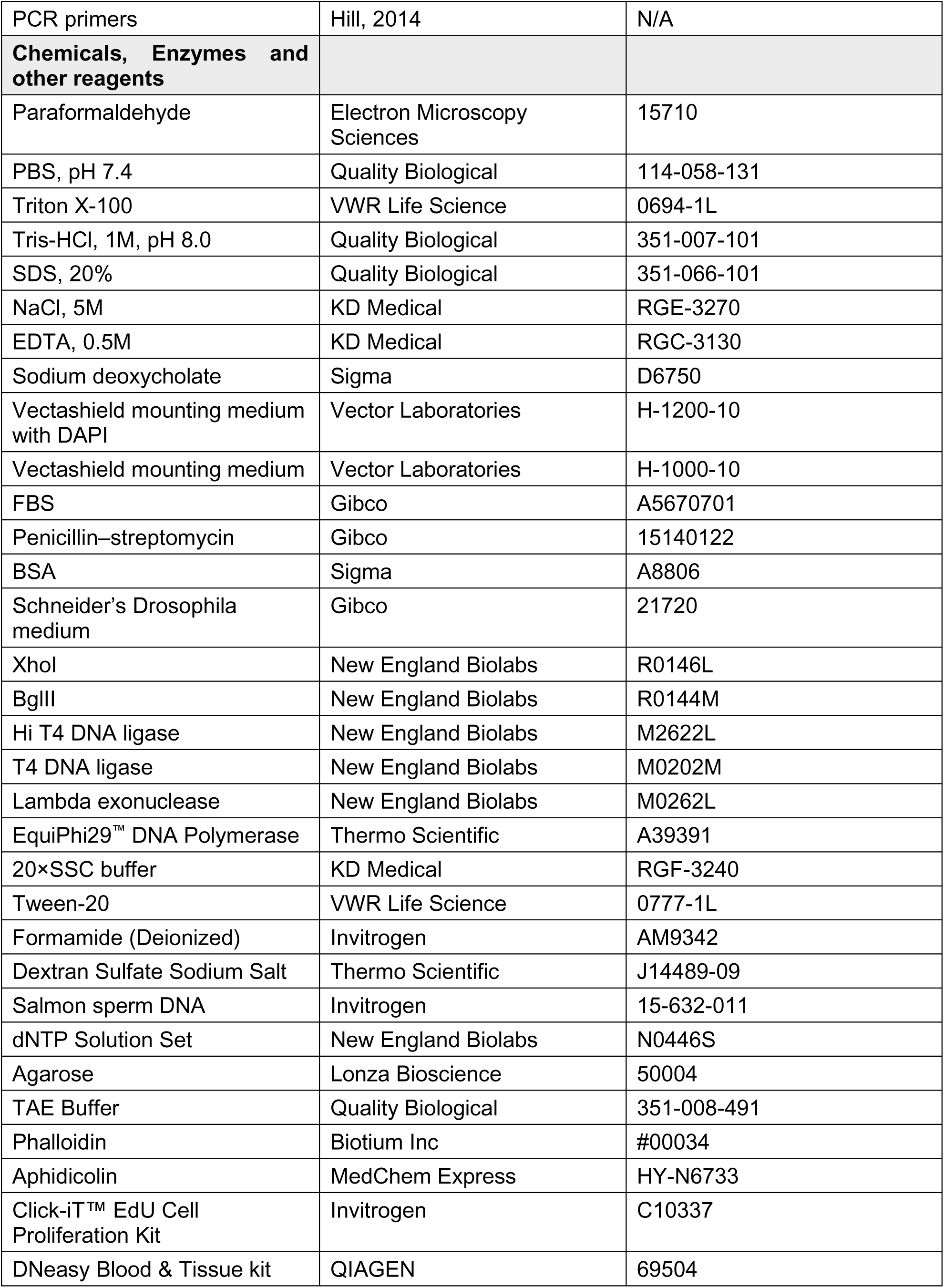
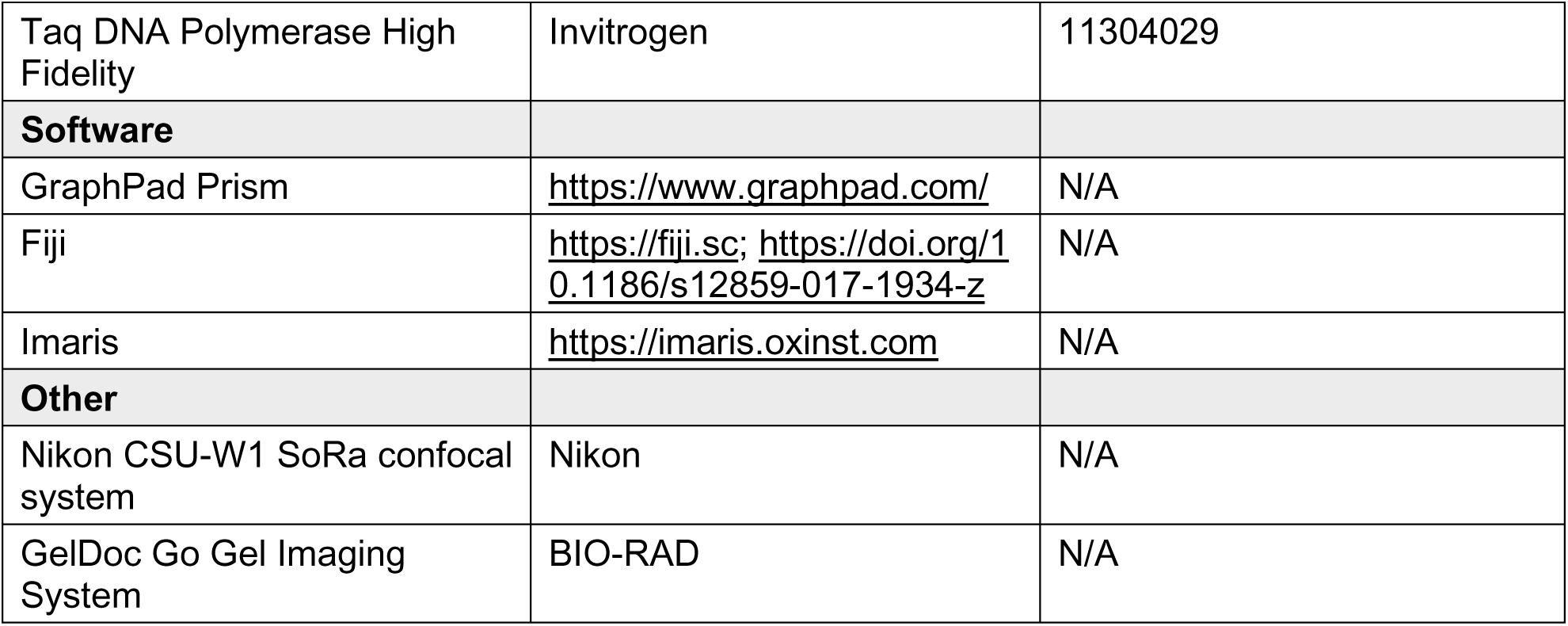
Reagents and Tools Table

**Supplementary Table 1.**
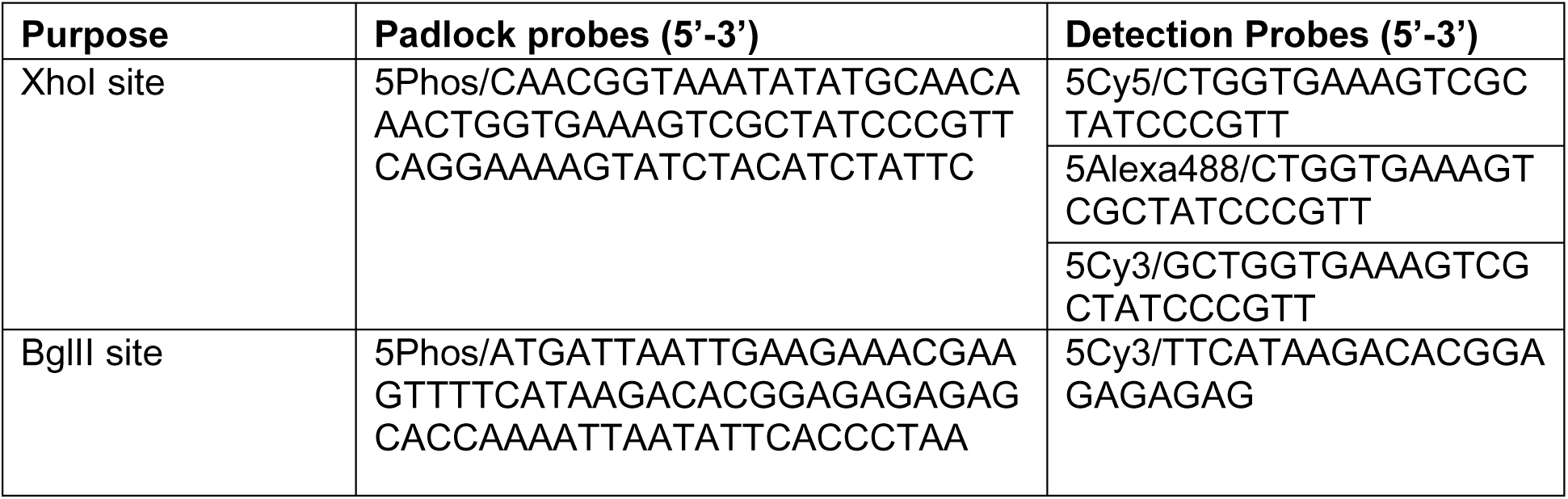

5Phos: a phosphorylation modification at the 5’ terminus of the oligonucleotide.

5Cy3: a Cy™3 dye modification at the 5’ terminus of the oligonucleotide.

5Cy5: a Cy™5 dye modification at the 5’ terminus of the oligonucleotide.

5Alexa488: an Alexa Fluor-488 dye modification at the 5’ terminus of an oligonucleotide.

The detection probes used for each experiment are as follows:

Figure 1C: Cy3-XhoI or Cy3-BglII.

Figure 2&3: Alexa488-XhoI and Cy3-BglII.

Figure 4: Cy5-XhoI and Cy3-BglII.

## Acknowledgments

This work was supported by the Intramural Research Program of the National Heart, Lung, and Blood Institute.

## Author contributions

C.Z. and H.X. conceived the project and designed the experiments. C.Z. performed the experiments. H.M. generated the heteroplasmic fly lines. C.Z., Z.C. analyzed the data. C.Z., Z.C., H.M., and H.X. wrote the manuscript.

## References

1. Bastock R, St Johnston D (2008) Drosophila oogenesis. Curr Biol 18: R1082–R1087

2. Birky CW (1994) Relaxed and Stringent Genomes - Why Cytoplasmic Genes Dont Obey Mendels Laws. J Hered 85: 355–365

3. Bogenhagen D, Clayton DA (1978) Mechanism of mitochondrial DNA replication in mouse L-cells: kinetics of synthesis and turnover of the initiation sequence. J Mol Biol 119: 49–68

4. Burman JL, Itsara LS, Kayser E-B Suthammarak W, Wang AM, Kaeberlein M, Sedensky MM, Morgan PG, Pallanck LJ (2014) A Drosophila model of mitochondrial disease caused by a complex I mutation that uncouples proton pumping from electron transfer. Dis Model Mech 7:1165–1174

5. Chandrasegaram R, Hynes-Allen AM, Gao B, Dhawanjewar A, Frison M, Petridi S, Chinnery PF, Ma H, van den Ameele J (2025) Single-molecule mitochondrial DNA imaging reveals heteroplasmy dynamics shaped by developmental bottlenecks and selection in different organs in vivo. bioRxiv: 2025.2001.2024.634671

6. Chen Z, Qi Y, French S, Zhang GF, Garcia RC, Balaban R, Xu H (2015) Genetic mosaic analysis of a deleterious mitochondrial DNA mutation in Drosophila reveals novel aspects of mitochondrial regulation and function. Mol Biol Cell 26: 674–684

7. Chen Z, Wang ZH, Zhang GF, Bleck CKE, Chung DJ, Madison GP, Lindberg E, Combs C, Balaban RS, Xu H (2020) Mitochondrial DNA segregation and replication restrict the transmission of detrimental mutation. J Cell Biol 219

8. Chen Z, Zhang F, Lee A, Yamine M, Wang ZH, Zhang GF, Combs C, Xu H (2025) Mitochondrial DNA removal is essential for sperm development and activity. Embo Journal

9. Chiang ACY, McCartney E, O’Farrell PH, Ma HS (2019) A Genome-wide Screen Reveals that Reducing Mitochondrial DNA Polymerase Can Promote Elimination of Deleterious Mitochondrial Mutations. Curr Biol 29: 4330–4336

10. Clary DO, Wolstenholme DR (1985) The mitochondrial DNA molecular of Drosophila yakuba: nucleotide sequence, gene organization, and genetic code. J Mol Evol 22: 252–271

11. Cree LM, Samuels DC, Lopes SCDS, Rajasimha HK, Wonnapinij P, Mann JR, Dahl HHM, Chinnery PF (2008) A reduction of mitochondrial DNA molecules during embryogenesis explains the rapid segregation of genotypes. Nat Genet 40: 249–254

12. de Cuevas M, Lilly MA, Spradling AC (1997) Germline cyst formation in Drosophila. Annu Rev Genet 31: 405–428

13. Fan WW, Waymire KG, Narula N, Li P, Rocher C, Coskun PE, Vannan MA, Narula J, MacGregor GR, Wallace DC (2008) A mouse model of mitochondrial disease reveals germline selection against severe mtDNA mutations. Science 319: 958–962

14. Floros VI, Pyle A, Dietmann S, Wei W, Tang WWC, Irie N, Payne B, Capalbo A, Noli L, Coxhead J et al (2018) Segregation of mitochondrial DNA heteroplasmy through a developmental genetic bottleneck in human embryos. Nat Cell Biol 20: 144–151

15. Gilkerson RW, Schon EA, Hernandez E, Davidson MM (2008) Mitochondrial nucleoids maintain genetic autonomy but allow for functional complementation. J Cell Biol 181: 1117–1128

16. Hill JH, Chen Z, Xu H (2014) Selective propagation of functional mitochondrial DNA during oogenesis restricts the transmission of a deleterious mitochondrial variant. Nat Genet 46: 389–392

17. Hinnant TD, Merkle JA, Ables ET (2020) Coordinating Proliferation, Polarity, and Cell Fate in the Female Germline. Front Cell Dev Biol 8

18. Isaac RS, Tullius TW, Hansen KG, Dubocanin D, Couvillion M, Stergachis AB, Churchman LS (2024) Single-nucleoid architecture reveals heterogeneous packaging of mitochondrial DNA. Nat Struct Mol Biol 31

19. Jenuth JP, Peterson AC, Fu K, Shoubridge EA (1996) Random genetic drift in the female germline explains the rapid segregation of mammalian mitochondrial DNA. Nat Genet 14: 146–151

20. Larsson C, Koch J, Nygren A, Janssen G, Raap AK, Landegren U, Nilsson M (2004) genotyping individual DNA molecules by target-primed rolling-circle amplification of padlock probes. Nat Methods 1: 227–232

21. Lieber T, Jeedigunta SP, Palozzi JM, Lehmann R, Hurd TR (2019) Mitochondrial fragmentation drives selective removal of deleterious mtDNA in the germline. Nature 570: 380–384

22. Liu Y, Chen Z, Wang ZH, Delaney KM, Tang JJ, Pirooznia M, Lee DY, Tunc I, Li YS, Xu H (2022) The PPR domain of mitochondrial RNA polymerase is an exoribonuclease required for mtDNA replication in. Nat Cell Biol 24: 757–765

23. Liu Y, Liu HB, Zhang F, Xu H (2024) The initiation of mitochondrial DNA replication. Biochem Soc T 52: 1243–1251

24. Ma H, O’Farrell PH (2015) Selections that isolate recombinant mitochondrial genomes in animals. Elife 4

25. Ma HS, Xu H, O’Farrell PH (2014) Transmission of mitochondrial mutations and action of purifying selection in. Nat Genet 46: 393–397

26. Merkle JA (2023) Dissection, Fixation, and Standard Staining of Adult Drosophila Ovaries. Methods Mol Biol 2626: 49–68

27. Palozzi JM, Jeedigunta SP, Minenkova AV, Monteiro VL, Thompson ZS, Lieber T, Hurd TR (2022) Mitochondrial DNA quality control in the female germline requires a unique programmed mitophagy. Cell Metab 34: 1809–1823 e1806

28. Pesole G, Gissi C, De Chirico A, Saccone C (1999) Nucleotide substitution rate of mammalian mitochondrial genomes. J Mol Evol 48: 427–434

29. Rand DM (2008) Mitigating mutational meltdown in mammalian mitochondria. PLoS Biol 6: e35

30. Ru YF, Deng XL, Chen JT, Zhang LP, Xu Z, Lv QY, Long SY, Huang ZJ, Kong MH, Guo J et al (2024) Maternal age enhances purifying selection on pathogenic mutations in complex I genes of mammalian mtDNA. Nature Aging 4

31. Spradling AC (1993) Germline Cysts - Communes That Work. Cell 72: 649–651

32. Stewart JB, Freyer C, Elson JL, Larsson NG (2008) Purifying selection of mtDNA and its implications for understanding evolution and mitochondrial disease. Nat Rev Genet 9: 657–662

33. Taylor RW, Turnbull DM (2005) Mitochondrial DNA mutations in human disease. Nat Rev Genet 6: 389–402

34. Wallace DC (2008) Anecdotal, historical and critical commentaries on genetics. Genetics 179: 727–735

35. Wang ZH, Liu Y, Chaitankar V, Pirooznia M, Xu H (2019) Electron transport chain biogenesis activated by a JNK-insulin-Myc relay primes mitochondrial inheritance in Drosophila. Elife 8

36. Wang ZH, Zhao WJ, Combs CA, Zhang F, Knutson JR, Lilly MA, Xu H (2023) Mechanical stimulation from the surrounding tissue activates mitochondrial energy metabolism in differentiating germ cells. Dev Cell 58: 2249–2260

37. Wei W, Tuna S, Keogh MJ, Smith KR, Aitman TJ, Beales PL, Bennett DL, Gale DP, Bitner-Glindzicz MAK, Black GC et al (2019) Germline selection shapes human mitochondrial DNA diversity. Science 364

38. Wolff JN, Sutovsky P, Ballard JW (2013) Mitochondrial DNA content of mature spermatozoa and oocytes in the genetic model Drosophila. Cell Tissue Res 353: 195–200

39. Xu H (2008) Manipulating the metazoan mitochondrial genome with targeted restriction enzymes. Science 321: 575–577

40. Yang Q, Liu P, Anderson NS, Shpilka T, Du Y, Naresh NU, Li R, Zhu LJ, Luk K, Lavelle J, Zeinert RD, Chien P, Wolfe SA, Haynes, CM (2022) LONP-1 and ATFS-1 sustain deleterious heteroplasmy by promoting mtDNA replication in dysfunctional mitochondria. Nat Cell Biol 24: 181–193

41. Zhang F, Lee A, Freitas AV, Herb JT, Wang ZH, Gupta S, Chen Z, Xu H (2024) A transcription network underlies the dual genomic coordination of mitochondrial biogenesis. Elife 13

42. Zhang KX, Deng RJ, Teng XC, Li Y, Sun YP, Ren XJ, Li JH (2018) Direct Visualization of Single-Nucleotide Variation in mtDNA Using a CRISPR/Cas9-Mediated Proximity Ligation Assay. J Am Chem Soc 140: 11293–11301

43. Zhang Y, Chen Y, Gucek M, Xu H (2016) The mitochondrial outer membrane protein MDI promotes local protein synthesis and mtDNA replication. EMBO J 35: 1045–1057

44. Zhang Y, Wang ZH, Liu Y, Chen Y, Sun N, Gucek M, Zhang F, Xu H (2019) PINK1 Inhibits Local Protein Synthesis to Limit Transmission of Deleterious Mitochondrial DNA Mutations. Mol Cell 73: 1127–1137

